# Assembly of SARS-CoV-2 ribonucleosomes by truncated N* variant of the nucleocapsid protein

**DOI:** 10.1101/2023.08.16.553581

**Authors:** Armin N. Adly, Maxine Bi, Christopher R. Carlson, Abdullah M. Syed, Alison Ciling, Jennifer A. Doudna, Yifan Cheng, David O. Morgan

## Abstract

The Nucleocapsid (N) protein of SARS-CoV-2 compacts the RNA genome into viral ribonucleoprotein (vRNP) complexes within virions. Assembly of vRNPs is inhibited by phosphorylation of the N protein SR region. Several SARS-CoV-2 variants of concern carry N protein mutations that reduce phosphorylation and enhance the efficiency of viral packaging. Variants of the dominant B.1.1 viral lineage also encode a truncated N protein, termed N* or Δ(1–209), that mediates genome packaging despite lacking the N-terminal RNA-binding domain and SR region. Here, we show that Δ(1–209) and viral RNA assemble into vRNPs that are remarkably similar in size and shape to those formed with full-length N protein. We show that assembly of Δ(1–209) vRNPs requires the leucine-rich helix (LH) of the central disordered region, and that the LH promotes N protein oligomerization. We also find that fusion of a phosphomimetic SR region to Δ(1–209) inhibits RNA binding and vRNP assembly. Our results provide new insights into the mechanisms by which RNA binding promotes N protein self-association and vRNP assembly, and how this process is modulated by SR phosphorylation.

## Introduction

Severe acute respiratory syndrome coronavirus 2 (SARS-CoV-2), the causative agent of the COVID-19 pandemic, is an enveloped betacoronavirus with a ∼30 kb single-stranded positive-sense RNA genome packaged inside a ∼100 nm virion (1–3). Compaction of the genome into the nucleocapsid depends on the nucleocapsid (N) protein, the most abundant viral protein in an infected cell (4–6).

Following infection and genome unpackaging, the first two-thirds of the SARS-CoV-2 genome is translated to produce numerous nonstructural proteins, including RNA-dependent RNA polymerase and other proteins responsible for rearranging membranes of the endoplasmic reticulum (ER) to assemble the replication-transcription complex (RTC). The RTC is a network of double-membrane vesicles that scaffold viral RNA synthesis and shield it from host innate immunity (7–10). The final third of the viral genome is then transcribed to produce short subgenomic RNAs encoding the four structural proteins that form the virion: spike (S), membrane (M), envelope (E), and nucleocapsid (N) (11–13). Subgenomic transcription involves a template-switching mechanism in which viral RNA polymerase transcribes a structural protein gene from the 3’ end of the genome. When it encounters a transcription-regulating sequence (TRS) at the end of a structural gene, the polymerase switches to a TRS located at the 5’ end of the genome to complete synthesis of a short negative-sense RNA, which is transcribed to a positive-sense RNA for translation (11,14). The S, M, and E proteins contain transmembrane domains that insert into the ER membrane at sites of viral assembly. The N protein accumulates in the cytoplasm at RTCs, where it likely plays a role in subgenomic transcription, and at nearby sites of viral assembly, where it packages the genome (7,8,15–22).

Cryo-electron tomography of intact SARS-CoV-2 virions has revealed that each virus contains 35 to 40 discrete viral ribonucleoprotein (vRNP) complexes, which are likely to be distributed along one copy of the genome (23,24). These vRNPs are ∼15 nm in diameter and are estimated to contain roughly 800 nt of RNA and 12 copies of N protein. When combined *in vitro*, purified N protein and viral RNA form 15 nm vRNP particles that closely resemble the vRNP structures observed within virions (25,26).

Assembly of vRNP complexes requires multiple N protein regions that mediate protein-RNA and protein-protein interactions (25,27–31). The 46 kDa N protein contains two globular RNA-binding domains, the N- and C-terminal domains (NTD and CTD), flanked by three regions of intrinsic disorder (Fig. 1A) (32–38). In solution, N protein exists predominantly as a dimer through a high-affinity dimerization interface on the CTD (38). N protein dimers can self-assemble into tetramers and higher-order oligomers that are likely to be important in vRNP assembly. Oligomerization depends on multiple N protein regions in the central and C-terminal disordered regions (33,37–45). Among these, a leucine-rich helix (LH, aa 215-234) in the central disordered linker was shown in recent biophysical studies to self-associate and promote N protein oligomerization (30,46). The stability of vRNP complexes depends in part on multivalent interactions between a long RNA and multiple N proteins (25,29,47). However, N protein oligomerization and vRNP assembly can be promoted *in vitro* with small RNA molecules that are unlikely to crosslink multiple N protein dimers, suggesting that vRNP formation depends, at least in part, on N-N interactions that are stimulated by RNA binding (25,26,30,31). Consistent with this possibility, RNA binding to the NTD induces conformational changes that enhance LH-mediated self-association of N protein (46,48).

**Figure 1.**
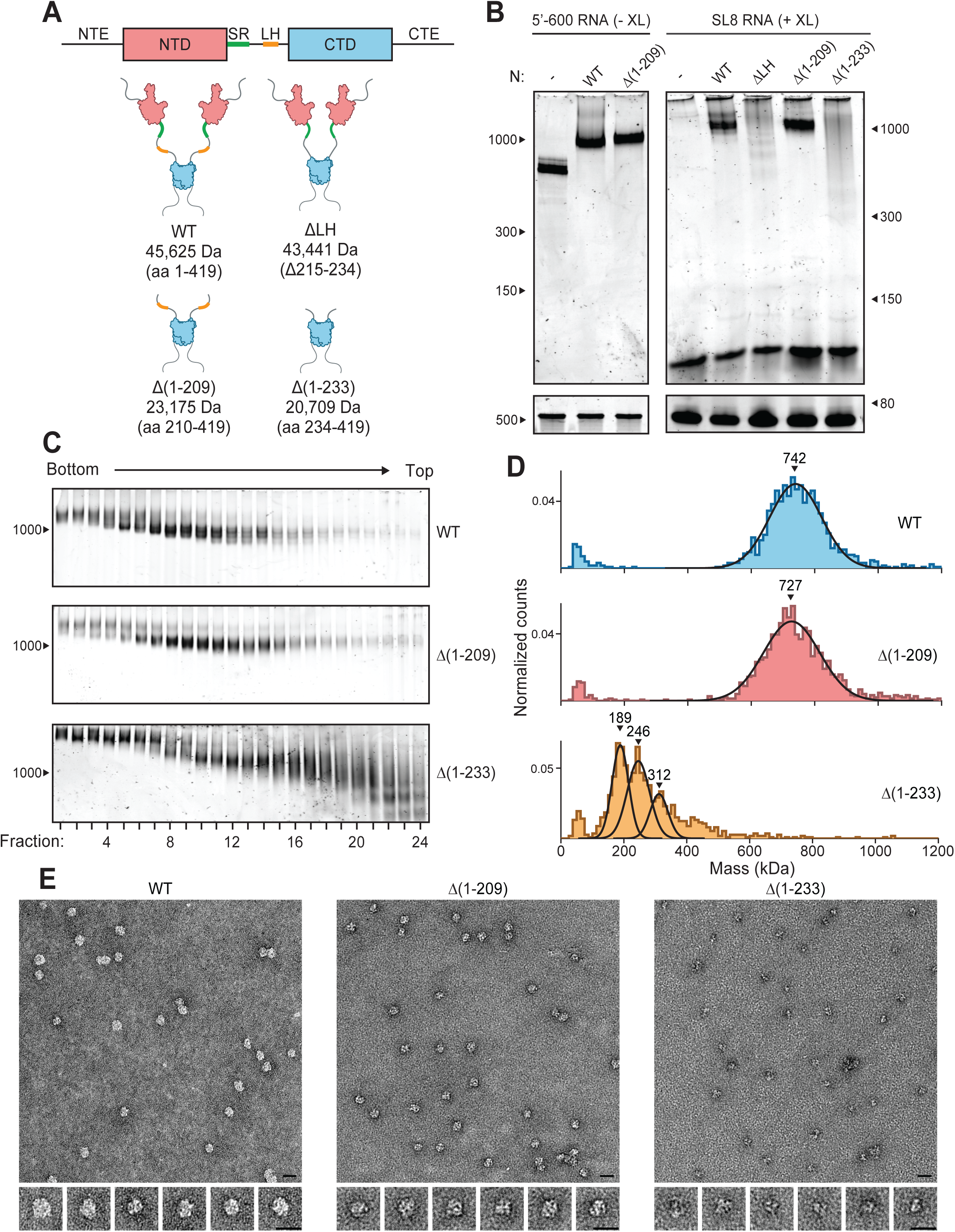
vRNPs formed by Δ(1-209) resemble those formed by full-length N protein. (A) Schematic of N protein domain architecture, including the N-terminal extension (NTE), N-terminal domain (NTD), serine/arginine region (SR), leucine-rich helix (LH), C-terminal domain (CTD), and C-terminal extension (CTE). Schematics comparing WT, ΔLH, Δ(1-209), and Δ(1-233) N protein mutants are also shown. Mass is that of monomeric N protein. (B) 15 μM N protein constructs were combined with 256 ng/μl 5’-600 RNA (left) or SL8 RNA (right) and analyzed by native (top) and denaturing (bottom) gel electrophoresis. SL8 ribonucleoprotein complexes were crosslinked (+XL) prior to native gel electrophoresis. Gels were stained with SYBR Gold to visualize RNA species. RNA length standards are shown on the left (nt). 256 ng/ul RNA was used to ensure all mixtures contain the same nucleotide concentration, regardless of RNA length. (C) Native gel analysis of vRNP complexes separated by glycerol gradient centrifugation in the presence of crosslinker (GraFix). 15 μM WT (top), Δ(1-209) (middle), or Δ(1–233) (bottom) was mixed with 256 ng/μl SL8 RNA, and added on top of a 10-40% glycerol gradient. 0.1% glutaraldehyde was added to the 40% glycerol buffer. RNA length standard shown on left (nt). (D) Selected fractions from the GraFix analyses in C were analyzed by mass photometry. Top, fractions 7+8 of WT in complex with SL8 RNA; middle, fractions 7+8 of Δ(1–209) in complex with SL8 RNA; bottom, fractions 22+23 of Δ(1–233) in complex with SL8 RNA. Data were fit to Gaussian distributions, with mean molecular mass indicated above each peak. Representative of at least two independent experiments (Table S1). The small peaks at the lower detection limit are likely to be artifacts seen in the analysis of GraFix fractions. (E) Negative stain electron microscopy of GraFix-purified vRNPs (combined fractions from C), with manually extracted images of single particles below. Left, WT in complex with SL8 RNA; middle, Δ(1–209) in complex with SL8 RNA; right, Δ(1–233) in complex with SL8 RNA. Scale bars are 20 nm. N, nucleocapsid; vRNP, viral ribonucleoprotein; WT, wild type.

The central disordered region also contains a conserved serine/arginine (SR)-rich sequence (aa 176-206), which contributes to RNA binding and is regulated by phosphorylation (4,21,27,49–51). The SR region is densely phosphorylated at multiple sites by host cell kinases in the cytoplasm but not inside the virion (21,52–56), implying that N is dephosphorylated at sites of virus assembly. Phosphorylation of N disrupts vRNP formation and appears to act as a structural switch to regulate the two major functions of N protein: unmodified N binds RNA to form compact vRNPs inside virions, whereas phosphorylated N maintains RNA in a less compacted state that may facilitate RNA transcription at the RTC (25,26,29,57–59).

Over the course of the pandemic, increasingly transmissible genetic variants of SARS-CoV-2 have emerged. Many of these variants carry N protein mutations that appear to improve viral packaging and infectivity, likely by enhancing the formation of vRNPs (60–64). All SARS-CoV-2 viral lineages contain independently arising mutations within the SR region of N protein, suggesting a strong link to viral fitness. Using a virus-like particle (VLP) system to assess the packaging of viral RNA by cultured cells, Syed et al. (61,63) showed that variant point mutations in the SR region increase production of VLPs while reducing phosphorylation of N. Treatment with kinase inhibitor also enhanced VLP assembly, suggesting that reduced phosphorylation is responsible for improved viral packaging. Conversely, reduction of phosphorylation by kinase inhibition or SR point mutation lessened RNA replication, supporting a role for phosphorylated N protein in efficient genome transcription (61). These results point to an adaptive conflict between the roles of phosphorylation in the two functions of N protein.

This adaptive conflict between assembly and replication might be resolved in part through the evolution of a truncated N protein termed N* or Δ(1–209), encoded by the B.1.1 lineage that includes the Alpha, Gamma, and Omicron variants (65–68). Mutations in this lineage create a new TRS that generates a short subgenomic RNA encoding a ∼23 kDa truncated version of N protein that is produced in addition to full-length N protein. Δ(1–209) is translated beginning at methionine 210 and therefore lacks the entire SR region and its phosphorylation sites, as well as the N-terminal RNA-binding domain (NTD), but retains the C-terminal dimerization and RNA-binding domain (CTD) (Fig. 1A). Recent work suggests that Δ(1–209) can package RNA and assemble infectious VLPs even in the presence of cellular kinase activity (61). These results suggest that a combination of Δ(1–209) and full-length N in infected cells provides a fitness benefit by allowing Δ(1–209) to function as a kinase-independent RNA packaging specialist in parallel with phosphorylated full-length N protein’s facilitation of RNA transcription.

In our previous work, we developed methods for the reconstitution of SARS-CoV-2 vRNPs from purified N protein and viral RNA fragments (25,26). Here, we applied these methods to test the hypothesis that the Δ(1–209) variant retains the RNA packaging function of full-length N protein. We show that Δ(1–209) and viral RNA assemble into vRNPs similar in size and shape to those formed with full-length N protein, despite lacking the NTD and SR region. We explore the biochemical properties, composition, and regulation of these vRNPs, providing new insights into the role of the disordered central linker region in packaging, and how its modification by phosphorylation regulates N function.

## Results

### vRNPs formed by Δ(1–209) resemble those formed by full-length N protein

We showed previously that full-length N protein and viral RNA form vRNP complexes *in vitro* that are similar in size and shape to vRNPs formed in virions (25,26). These vRNPs are estimated to contain 12 copies of N protein, or 6 N protein dimers. Assembly of vRNPs depends on multiple regions of N protein that promote numerous protein-protein and protein-RNA interactions.

To test the ability of Δ(1–209) to form vRNPs *in vitro*, we mixed purified N protein (Fig. S1) with a 600-nt RNA fragment from the 5’ end of the genome (5’-600) and analyzed the resulting RNA-protein complexes by gel electrophoresis on a native TBE gel (Fig. 1B, left). Both full-length N (WT) and Δ(1–209) bound RNA, forming a large species near the 1000-nt marker, suggesting that the truncated N protein was assembled into vRNPs.

In our previous work, we showed that vRNP assembly can also be achieved using a short, 72-nt stem-loop (termed SL8) from within the 5’-600 region (25). Unlike the large 600-nt RNA, which can serve as a scaffold to bind multiple copies of N protein, the short SL8 stem-loop RNA does not seem to engage in multivalent protein interactions, resulting in a less stable vRNP that is primarily dependent on protein-protein interactions between N proteins. We mixed WT or Δ(1–209) protein with SL8, crosslinked with 0.1% glutaraldehyde to stabilize the resulting complexes, and assessed vRNP formation by native gel electrophoresis (Fig. 1B, right). WT and Δ(1–209) bound SL8 RNA to produce vRNP complexes of similar sizes.

vRNP complexes were purified for further analysis by velocity sedimentation on a glycerol gradient as described previously (25,69). WT or Δ(1–209) were mixed with SL8 RNA and separated by centrifugation on a 10–40% glycerol gradient containing 0.1% glutaraldehyde crosslinker to stabilize the RNA-protein complex during centrifugation (a technique known as gradient fixation, or GraFix) (70). Individual fractions were then analyzed by native gel electrophoresis (Fig. 1C). Both WT and Δ(1–209) generated a broad vRNP peak migrating at the same size (at the 1000-nt marker) as in gel electrophoresis of unpurified samples. Peak fractions (7 and 8) of purified complexes were pooled and analyzed by mass photometry (Fig. 1D; note that our mass photometry figures provide representative examples, but we cite in the text the mean mass ± SD from 2 or 3 experiments listed in Table S1). For WT, we observed a broad peak centered at 736 ± 7 kDa, representing vRNP complexes with a likely stoichiometry of 6 N protein dimers bound to approximately 8 stem-loop RNAs (predicted mass of SL8: 23.1 kDa; predicted mass of WT dimer: 91.2 kDa; total predicted mass: 732.3 kDa). For Δ(1–209), we observed a broad peak at 727 ± 4 kDa, indicating that vRNP complexes assembled by the truncated N variant are remarkably similar in mass to the vRNP complexes formed by WT protein, despite the loss of half of the protein’s mass. These results might suggest that Δ(1–209) vRNPs incorporate roughly twice as many N proteins to form a similarly sized complex (predicted mass of Δ(1–209) dimer: 46.4 kDa).

As in our previous work, negative stain electron microscopy (EM) of the GraFix-purified WT vRNP revealed discrete heterogeneous 15 nm particles (Fig. 1E, left) that are similar in shape and size to vRNP complexes observed within virions by cryo-electron tomography (23–26). Negative stain EM of GraFix-purified Δ(1–209) revealed discrete structures that closely resemble vRNPs assembled with WT protein. Together with the mass photometry data, these surprising results indicate that the Δ(1–209) protein assembles vRNPs that are wild-type in size and general morphology despite major differences in the size of N protein and the likely number of N proteins in the complex (Fig. 1E, right). The RNA-binding NTD and SR region are not required for vRNP formation by Δ(1–209), suggesting that the RNA-binding groove on the CTD is sufficient for vRNP assembly.

### The leucine-rich helix is critical for Δ(1–209) vRNP assembly

Recently, Zhao et al. (30,46) demonstrated that the LH of the central disordered linker can self-assemble into trimers and tetramers that promote the formation of higher-order N protein complexes. In our previous work, we showed that deletion of the LH disrupts vRNP assembly *in vitro* (25). Here, we addressed the role of the LH in assembly of vRNPs formed by the Δ(1–209) protein, whose sequence begins just before the LH (Fig. 1A). We removed the LH to create an additional deletion mutant, Δ(1–233).

Mutant N proteins (Fig. S1) were mixed with SL8 RNA, crosslinked, and analyzed by native gel electrophoresis (Fig. 1B, right). As seen previously, deletion of the LH in full-length N protein disrupted vRNP assembly, resulting in a laddering of ribonucleoprotein subcomplexes on the gel. The Δ(1–233) mutant also displayed a major defect in vRNP assembly, suggesting that the LH is also required for the formation of vRNPs with Δ(1–209).

GraFix purification of the Δ(1–233) mutant in complex with SL8 RNA revealed a clear shift toward lower molecular mass species when compared to Δ(1–209), suggesting that Δ(1–233) vRNPs were not fully assembled and contained subcomplexes (Fig. 1C). Mass photometry analysis of fractions 22+23 from Δ(1–233) confirmed the presence of subcomplexes (Fig. 1D; Table S1). Δ(1–233) formed major species at 188 ± 2 kDa, 242 ± 6 kDa, and 305 ± 11 kDa. 188 kDa is the mass expected for three N dimers bound to three SL8 RNAs (predicted mass of SL8: 23.1 kDa; predicted mass of Δ(1–233) dimer: 41.4 kDa; predicted mass of complex: 193.5 kDa), and the stepwise ∼50-60 kDa increases in molecular mass are consistent with the addition of an N dimer bound to one SL8 RNA. Negative Stain EM of the GraFix-purified Δ(1–233) complex (fractions 22+23) revealed a smaller overall structure that is distinct from that of fully formed vRNP complexes assembled with WT or Δ(1–209) (Fig. 1E). These data suggest that protein-protein interactions mediated by the LH promote the formation of stable vRNPs by both the WT and Δ(1–209) proteins.

### The leucine-rich helix mediates N protein oligomerization

In the absence of RNA, N protein self-assembles into higher order oligomers through low-affinity protein-protein interactions that depend on the LH and other regions (30,31,38–40,42,46,71). RNA binding is thought to stimulate these interactions (see Introduction). We used mass photometry to explore RNA-independent N protein self-assembly by the truncated Δ(1–209) protein (Figs. 2A, 2B, and S2; Table S1). This method requires dilution to low protein concentrations (∼100 nM), where oligomerization of WT N protein is minimal. To capture the weak protein-protein interactions contributing to N protein oligomerization, samples were crosslinked with 0.1% glutaraldehyde before dilution.

**Figure 2.**
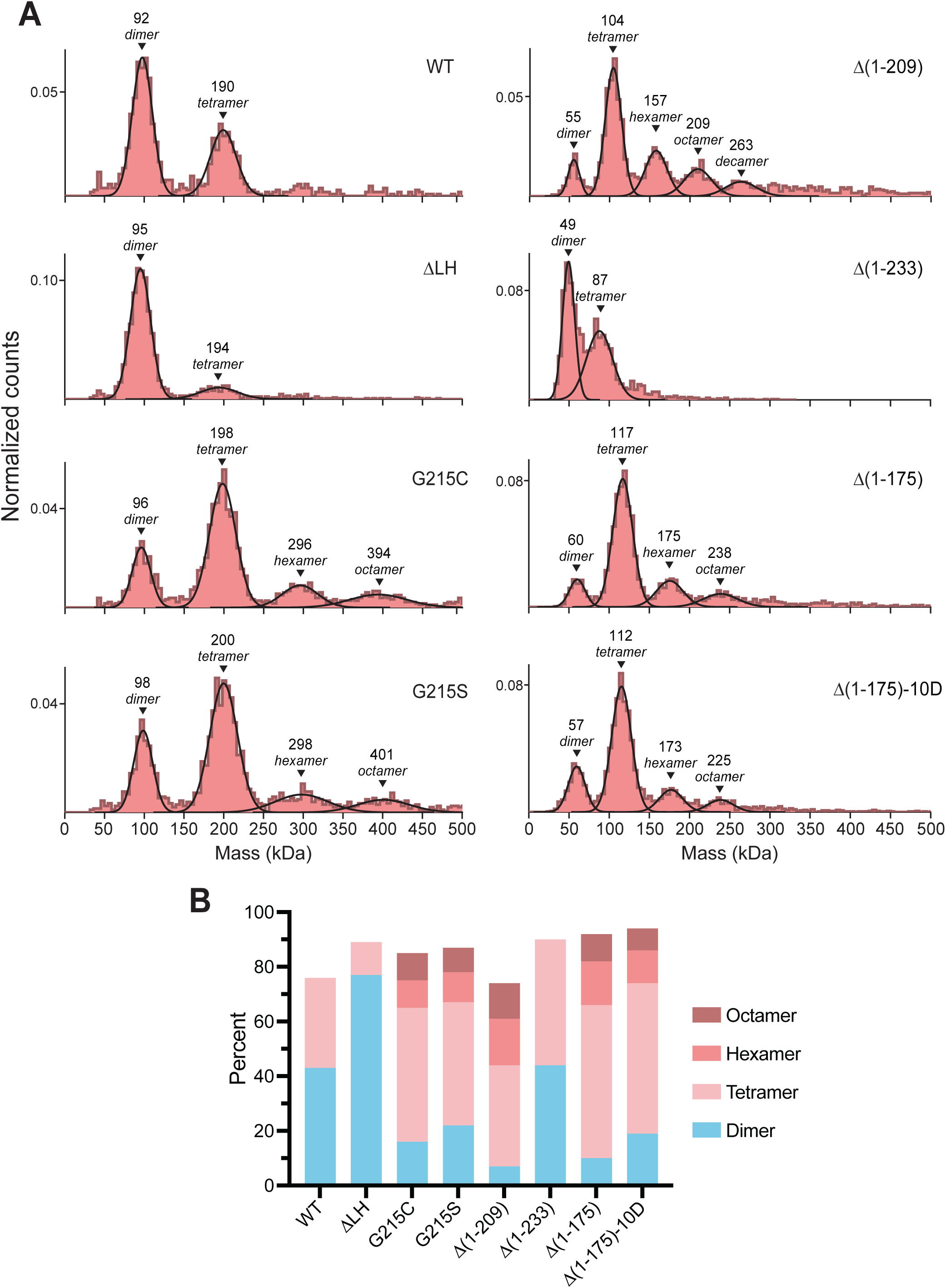
The leucine-rich helix mediates N protein oligomerization. (A) Mass photometry analysis of indicated N protein mutants in the presence of crosslinker. N protein (15 μM) was crosslinked with 0.1% glutaraldehyde prior to dilution and analysis. Data were fit to Gaussian distributions, with mean molecular mass and corresponding oligomeric state indicated above each peak. Representative of two independent experiments (Table S1). Parallel analysis of non-crosslinked samples in Fig. S2. (B) Mass photometry counts (as a percentage of total signal) were calculated for each detected oligomeric species in the indicated N protein mutants. Total percent does not equal 100 due to background signals at other masses. Means calculated from two independent experiments (Table S1). LH, leucine helix; N, nucleocapsid; WT, wild type.

For WT N protein, the majority of molecules were found in a dimer at 95 ± 4 kDa (predicted mass: 91.3 kDa), with a small population existing as tetramers at 195 ± 6 kDa (predicted mass: 182.5 kDa). Deletion of the LH (ΔLH) significantly reduced the tetramer population, confirming that the LH plays a critical role in RNA-independent N protein oligomerization.

Some SARS-CoV-2 variants have evolved mutations in N protein that enhance the protein-protein interactions mediating vRNP assembly (30,46,51). An excellent example is the G215C point mutation found in the delta variant, in which the glycine immediately N-terminal of the LH is substituted with a cysteine. This mutation induces conformational changes in the LH that enhance N protein oligomerization (30,46). These structural changes occur even in the absence of covalent disulfide bonds, as replacing the cysteine with a serine (G215S) has a similar effect. Mass photometry revealed that both the G215C and G215S mutations enhanced oligomerization (Figs. 2A, 2B, and S2; Table S1). Compared to WT, the G215C and G215S mutations shifted the predominant oligomer from dimer to tetramer, with additional smaller populations assembling into hexamers and octamers.

Analysis of crosslinked Δ(1–209) protein revealed a heterogeneous mixture dominated by tetramers (observed mass: 105 ± 1 kDa; predicted mass: 92.8 kDa), with hexamers, octamers, and decamers also appearing as distinct peaks. In contrast, the mass distribution for Δ(1–233) showed a significant reduction in the tetramer population in favor of dimers, once again pointing to the critical role of the LH in N protein oligomerization.

The enhanced oligomerization of Δ(1–209) compared to WT suggests that deletion of the NTD and SR region improves N protein self-association in the absence of RNA. This finding raises the possibility that the NTD acts as an antagonist to LH self-association, and deletion of the NTD in Δ(1–209) frees the LH to self-associate (see Discussion).

### Fusion of phosphomimetic SR region disrupts vRNP assembly by Δ(1–209)

In our previous work (25,26), we observed defects in vRNP formation with a phosphomimetic full-length N protein (the 10D mutant, in which 10 serines and threonines in the SR region are replaced with aspartic acid), as well as with N protein that had been phosphorylated *in vitro*. We found that phosphomimetic and phosphorylated N proteins form elongated, heterogeneous vRNPs, maintaining RNA in an uncompacted state that might support N protein’s role in viral transcription.

To provide further mechanistic insight into the function of N protein phosphorylation and dephosphorylation during coronavirus infection, we added the SR region or phosphomimetic SR region to the N-terminus of Δ(1–209), creating the Δ(1–175) and Δ(1–175)-10D proteins (Figs. 3A and S1). We mixed SL8 RNA with Δ(1–175) or Δ(1–175)-10D protein, crosslinked the resulting complexes, and analyzed vRNP formation by native gel electrophoresis (Fig. 3B). Δ(1–175) vRNPs shifted to a larger species than Δ(1–209) vRNPs, potentially representing an over-assembled vRNP complex containing more N protein dimers bound to additional RNAs. Native gel analysis also revealed a decrease in free SL8 RNA in the Δ(1–175) sample compared to Δ(1–209), corroborating SR involvement in RNA binding. Δ(1–175)-10D had a major defect in vRNP assembly with the SL8 RNA, resulting in a smeared band and the disappearance of a sharp, discrete vRNP band on native gel electrophoresis.

**Figure 3.**
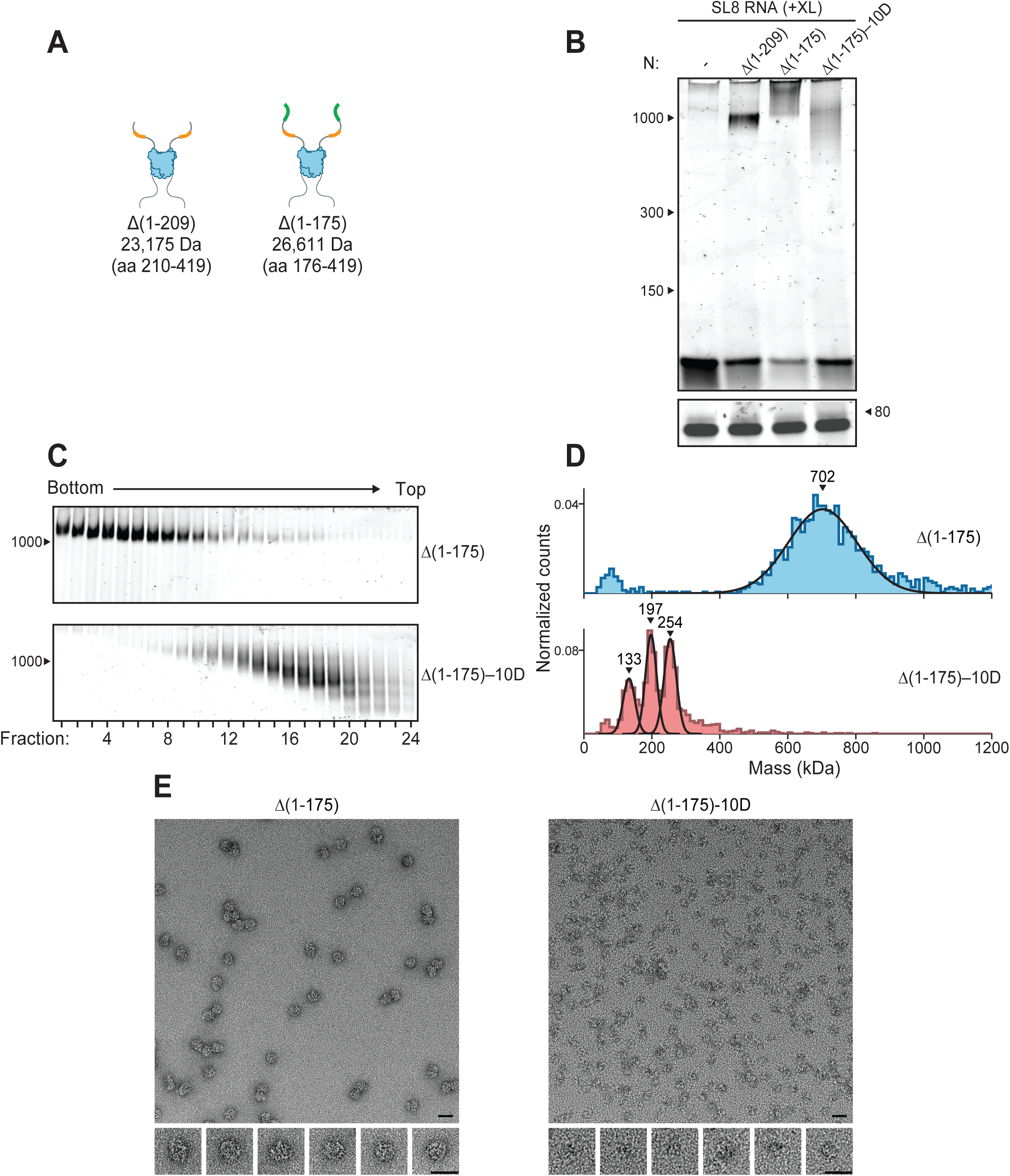
Fusion of phosphomimetic SR region to Δ(1-209) disrupts vRNP assembly. (A) Schematic comparing Δ(1-209) and Δ(1-175) proteins. Mass is that of monomeric N protein. (B) 15 μM N protein mutants were mixed with 256 ng/μl SL8 RNA, crosslinked (XL), and analyzed by native (top) and denaturing (bottom) gel electrophoresis. RNA length standards shown on left (nt). (C) 15 μM Δ(1-175) (top) or Δ(1-175)-10D (bottom) was mixed with 256 ng/μl SL8 RNA and separated by glycerol gradient centrifugation in the presence of crosslinker (GraFix). Fractions were collected and analyzed by native gel electrophoresis. RNA length standard shown on left (nt). (D) Selected fractions from the GraFix analyses in C were analyzed by mass photometry. Top: fractions 7+8 of Δ(1-175); bottom: fractions 19-22 of Δ(1-175)-10D. Representative of two independent experiments (Table S1). (E) Negative stain electron microscopy of GraFix-purified vRNPs from C, with manually extracted images of single particles below. Left, fractions 7+8 of Δ(1-175) in complex with SL8 RNA; right, fractions 19-22 of Δ(1-175)-10D in complex with SL8 RNA. Scale bars are 20 nm. N, nucleocapsid; vRNP, viral ribonucleoprotein.

The GraFix-purified Δ(1–175) complex migrated as a uniformly high molecular mass species (Fig. 3C, top). Analysis of peak fractions (7+8) by mass photometry indicated that these vRNPs were similar in mass to vRNPs assembled with Δ(1–209) (705 ± 4 kDa) (Fig. 3D, top; Table S1). Purification of the Δ(1–175)-10D complex by GraFix revealed a clear shift toward lower molecular mass species, with lower bands representing subcomplexes (Fig. 3C, bottom). Mass photometry of fractions 19-22 further confirmed the defect in vRNP assembly with Δ(1–175)-10D (Fig. 3D, bottom), as indicated by a series of small peaks separated by the combined mass of a single N dimer and RNA molecule. Negative stain EM of the GraFix-purified Δ(1–175) sample revealed discrete particles similar to vRNPs assembled with Δ(1–209). In contrast, the Δ(1–175)-10D mutant formed poorly defined complexes with a smaller, more diffuse structure (Fig. 3E), confirming the inability of the 10D phosphomimetic mutant to form stable, compact vRNPs. These data show that the addition of a phosphomimetic SR region to the N-terminus of Δ(1–209) inhibits vRNP formation.

We tested the possibility that the phosphomimetic mutations in Δ(1–175)-10D inhibit vRNP formation by disrupting LH-mediated oligomerization of N protein. Mass photometry revealed that multimerization of the Δ(1–175) and Δ(1–175)-10D proteins was similar to that seen with Δ(1–209) (Figs. 2A, 2B, and S2; Table S1). Thus, the phosphomimetic mutation, and presumably phosphorylation, does not have a major impact on RNA-independent N protein interactions that promote vRNP formation.

### Phosphorylation of SR region inhibits RNA binding

The SR region has been implicated in N protein oligomerization and RNA binding (27,42,44,49–51). In our previous work, we found that deletion of the SR region in full-length N protein prevented vRNP formation (25). We also observed that the 10D phosphomimetic mutation had a greater effect on the behavior of RNA-dependent condensates than an SR deletion, suggesting that phosphorylation does not simply inhibit the function of the SR region but has additional antagonistic effects (26). We hypothesize that the dense array of negative charges on the phosphorylated (or phosphomimetic) SR region blocks RNA binding to the SR region and also creates a disordered polyanion that binds to positively-charged RNA-binding sites on the protein, thereby interfering with RNA binding and vRNP formation. In the case of Δ(1–209), the major RNA binding site is on the CTD.

To investigate the effects of phosphomimetic mutations on RNA binding, we used fluorescence polarization to measure equilibrium binding of a fluorescent 10-nt RNA (of random sequence) to N protein mutants (Fig. 4A). Δ(1–209) bound the RNA with a dissociation constant K_D_ of 468 ± 41 nM, whereas Δ(1–175) N had a K_D_ of 114.0 ± 9 nM. The addition of the SR region therefore caused a ∼4-fold increase in affinity, consistent with previous evidence that the SR region contributes to RNA binding. In contrast, Δ(1–175)-10D bound with a K_D_ of 1049 ± 52 nM, representing a ∼2-fold decrease in affinity for RNA relative to the Δ(1–209) protein. These results are consistent with the idea that phosphorylation blocks RNA binding to the SR region but also has additional inhibitory effects on RNA binding at other sites.

**Figure 4.**
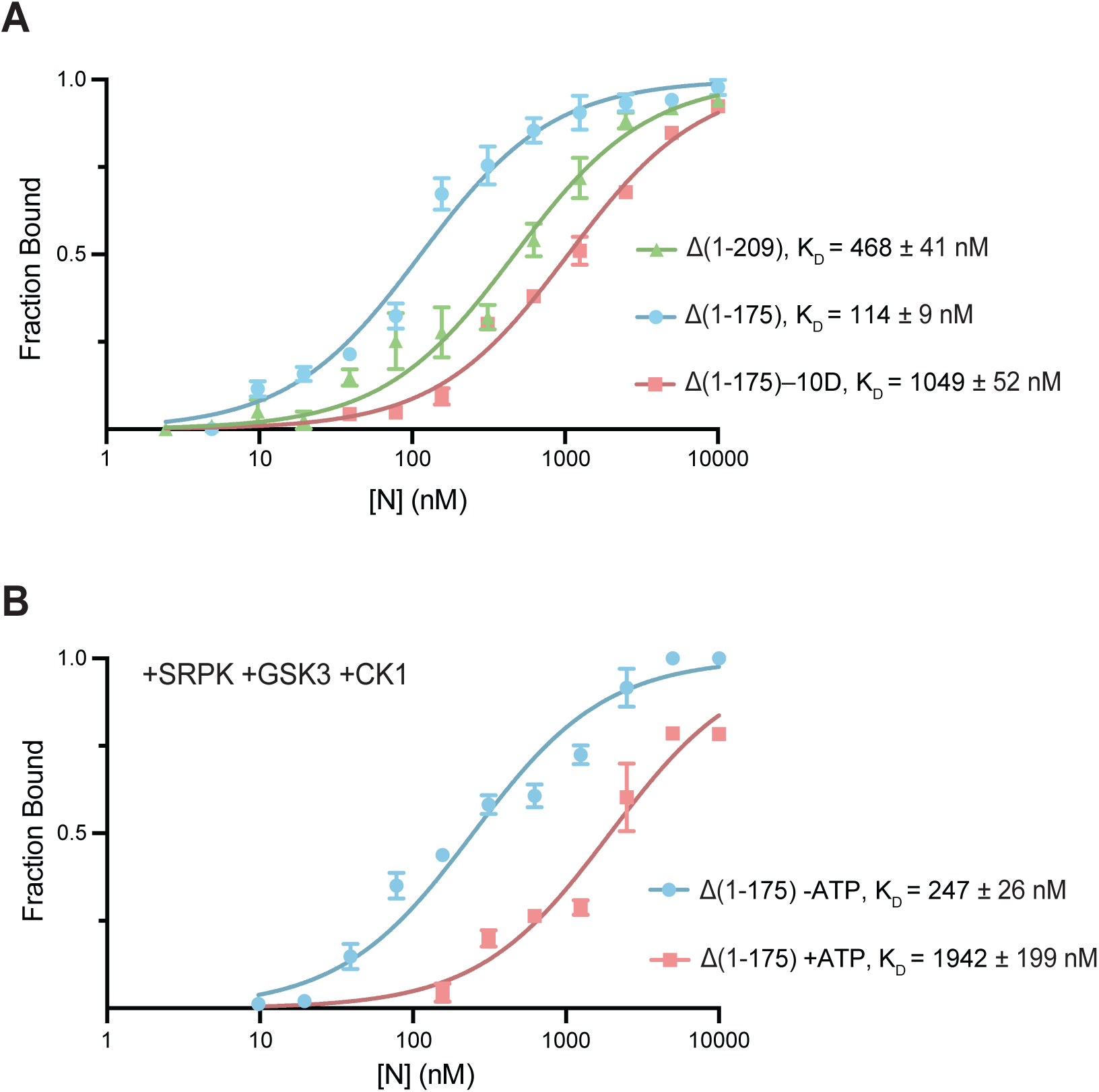
Phosphomimetic SR region competitively inhibits RNA binding. (A) Fluorescence polarization of 10-nt RNA binding to Δ(1-209), Δ(1-175), or Δ(1-175)-10D protein. Indicated concentrations of N protein mutants were incubated with 10 nM N_10_ RNA (a degenerate 10-nt RNA oligo with a 3’-FAM modification) and fluorescence polarization was recorded. Data points represent mean ± SEM of three independent experiments. K_D_ of each mutant is shown. (B) Fluorescence polarization of 10-nt RNA binding to Δ(1-175) protein phosphorylated *in vitro*. Δ(1-175) protein was incubated with the indicated kinases in the absence (blue) or presence (red) of ATP, mixed with 10 nM N_10_ RNA, and analyzed by fluorescence polarization. Data points represent mean ± SEM of three independent experiments. K_D_ of each mutant is shown. CK1, casein kinase 1; GSK3, glycogen-synthase kinase 3; N, nucleocapsid; SRPK, serine-arginine protein kinase.

We also tested RNA binding affinity with Δ(1–175) N protein that had been phosphorylated *in vitro* by SRPK, GSK3, and CK1 (Fig. 4B), the kinases implicated in the sequential phosphorylation of N protein (55). Phosphorylated Δ(1–175) bound the RNA with a K_D_ of 1942 ± 199 nM, representing an ∼8-fold lower affinity compared to unphosphorylated N (K_D_ = 247 ± 26 nM). These results suggest that a phosphorylated SR region disrupts vRNP assembly by inhibiting protein-RNA interactions.

### The leucine-rich helix is critical for virus-like particle assembly by Δ(1–209)

To compare the ability of N protein mutants to support viral assembly in live cells, we used a recently developed method for the production of virus-like particles (VLPs) by expression of the four SARS-CoV-2 structural proteins plus a luciferase reporter RNA (Figs. 5A, 5B) (61–63). Luminescence is used as a measure of luciferase mRNA expression in receiver cells, thereby serving as a marker of the efficiency of VLP packaging and assembly.

**Figure 5.**
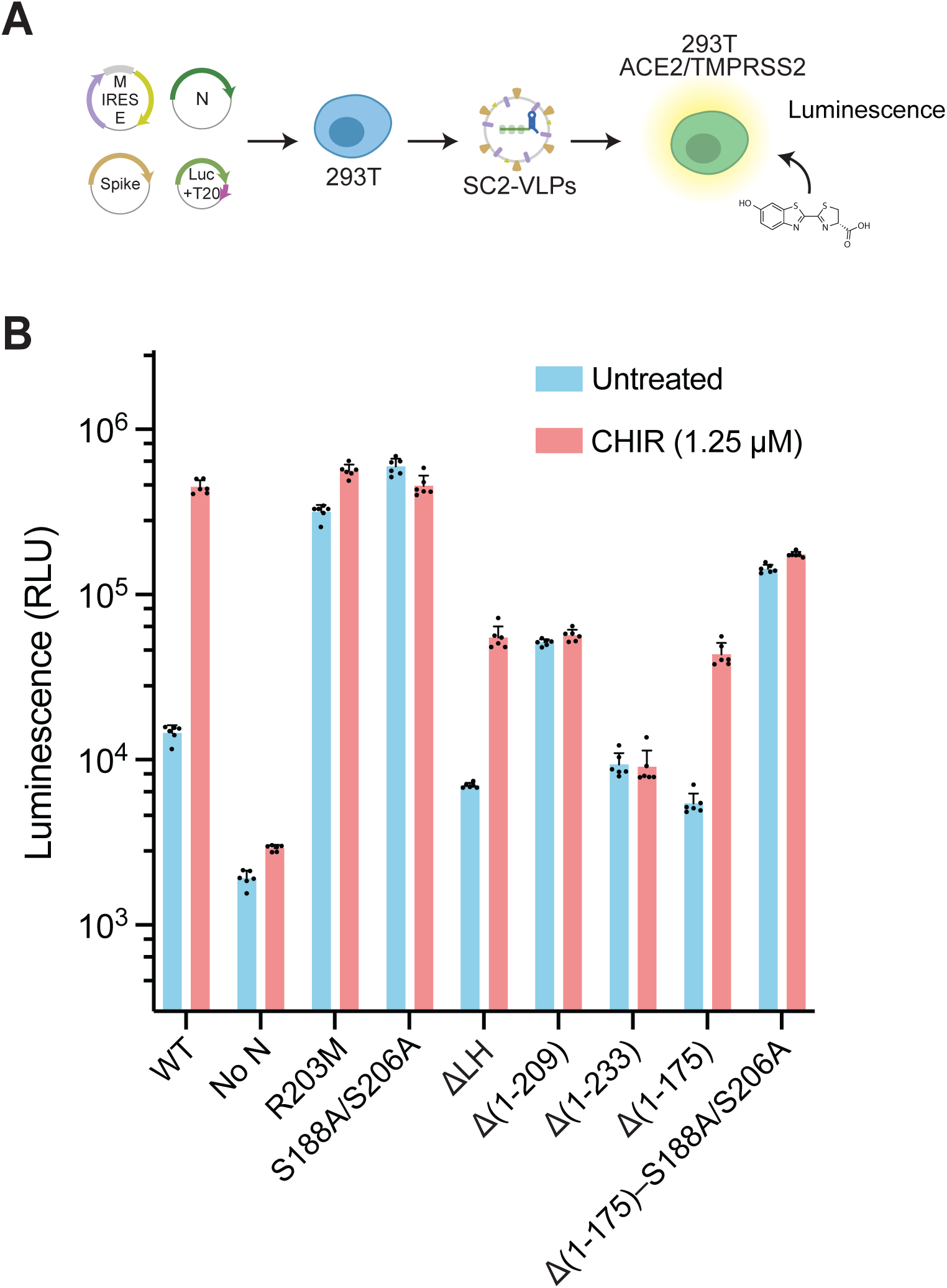
The leucine-rich helix is critical to virus-like particle assembly by Δ(1-209). (A) Schematic of the process for production and analysis of SARS-CoV-2 virus-like particles (SC2-VLPs). VLPs carrying luciferase (Luc) and packaging (T20) RNA were generated in 293T cells expressing wild-type (WT) or variant N protein. VLPs were added to 293T receiver cells expressing viral entry factors ACE2 and TMPRSS2, and luminescence due to luciferase expression was measured. (B) VLPs were generated with the indicated N protein, in the absence (blue) or presence (red) of 1.25 μM CHIR 98014, a GSK3 inhibitor. Data points represent mean (± SD) of 6 independent transfections. E, envelope; GSK3, glycogen-synthase kinase 3; LH, leucine helix; M, membrane; N, nucleocapsid; RLU, relative luminescence units.

The VLP system allows analysis of the impact of phosphorylation on the packaging function of N protein. The SR region of N is phosphorylated by a kinase cascade beginning with SRPK-mediated phosphorylation at S188 and S206, which primes N protein for sequential phosphorylation by GSK3 and CK1, resulting in a potential total of 14 phosphorylated residues (4,21,25,26,52,54–56). Treatment of VLP-producing cells with GSK3 inhibitor (CHIR98014) allows analysis of the effects of phosphorylation on VLP production.

As seen previously (61), treatment with GSK3 inhibitor greatly enhanced VLP assembly by wild-type N protein (Fig. 5B). Similarly, SR mutations that reduce phosphorylation (R203M, S188A/S206A) enhanced VLP assembly, and packaging by these mutants was unaffected by kinase inhibition.

The truncated Δ(1–209) protein was also capable of packaging RNA and producing infectious VLPs, as seen recently (61). Δ(1–209) lacks the phosphorylated SR region, and thus its packaging function was not affected by GSK3 inhibition.

Deletion of the LH from full-length N protein reduced VLP assembly ∼2-fold in untreated cells and ∼8-fold in cells treated with kinase inhibitor. Similarly, Δ(1–233) showed ∼6-fold reduced VLP assembly relative to Δ(1–209) in both untreated and CHIR98014-treated cells. These data suggest that the LH is important for viral assembly and RNA packaging, consistent with evidence *in vitro* that removal of the LH weakens N protein self-association and disrupts vRNP formation.

Δ(1–175) displayed ∼9-fold lower packaging than Δ(1–209) in the presence of kinases, consistent with our finding that phosphomimetic mutations of the SR region disrupt vRNP formation *in vitro*. Treatment with kinase inhibitor resulted in the expected increase in packaging, boosting Δ(1–175) VLP assembly to levels comparable to those of Δ(1–209). Δ(1–175)-S188A/S206A produced ∼3-fold higher VLP assembly than Δ(1–209) in both untreated and CHIR-98014-treated cells. This suggests that the unphosphorylated SR region promotes packaging, consistent with evidence that the SR region is an RNA-binding site that helps drive vRNP formation.

## Discussion

Our findings provide new mechanistic insight into the current model for SARS-CoV-2 vRNP assembly and its phosphoregulation. We show that the truncated Δ(1–209) protein from the B.1.1 lineage assembles vRNPs similar in size and shape to those formed with full-length N. Assembly of these vRNPs depends on the self-associating leucine-rich helix of the central disordered linker. Surprisingly, however, assembly does not require the major RNA-binding sites of the NTD and SR region, leaving only the CTD as the sole major RNA binding site involved in vRNP assembly by Δ(1–209).

The precise structure of the SARS-CoV-2 ribonucleosome remains mysterious. Our previous work (25) suggested that six N protein dimers organize several hundred bases of RNA, much of which is folded into a heterogeneous array of double-stranded stem-loops and other secondary structures (72,73). Although our reconstituted vRNPs appear uniform in diameter and general morphology, our analyses by negative-stain EM and cryo-electron microscopy suggest that these vRNPs are highly heterogeneous, precluding high-resolution structural analyses. We speculate that the densities in our EM images represent flexible RNA stem-loops organized by a dynamic protein framework held together by low-affinity protein-protein interactions. Our new evidence that the Δ(1–209) protein organizes vRNPs of normal size provides new clues and new questions. The vRNP appears to maintain its size when one of the two major RNA-binding sites of the N protein is missing, suggesting that the number of N proteins in the vRNP is increased, by unknown mechanisms, to provide additional RNA-binding sites.

Previous evidence suggests that the binding of small RNA molecules induces a conformational change in N protein that promotes oligomerization and initiates the assembly of higher-order vRNP complexes (30,31). In an NMR study, Pontoriero et al. (48) found that the leucine-rich helix undergoes a resonance shift in response to short stem-loop RNA binding to the NTD, suggesting a regulatory mechanism by which intramolecular interactions between the NTD and LH are disrupted by RNA binding. In other recent studies, Zhao et al. (46) similarly found that small oligonucleotide binding induces conformational changes that enhance N protein self-association at the LH. Here, we found that Δ(1–209) is capable of forming vRNPs despite lacking the NTD, and that Δ(1–209) exhibits improved self-association compared to full-length N. We speculate that RNA binding to the NTD promotes N protein self-association and initiates vRNP assembly by releasing the LH from an inhibitory interaction with the NTD. Given that Δ(1–209) doesn’t require the NTD to form vRNPs, the NTD and SR may serve as a regulatory switch that unleashes the LH to promote RNA-dependent oligomerization.

The SR region contributes to vRNP formation, probably by serving as an additional RNA binding site adjacent to the NTD. The presence of the SR region in the Δ(1–175)-S188A/S206A mutant enhances VLP assembly when compared to Δ(1–209), suggesting that the SR region is important for packaging. In our previous work, we showed the SR region is required for vRNP formation with full-length N, as deletion of the SR or replacement of the SR region with a random linker caused defects in vRNP formation (25). According to the model described above, deletion of both the NTD and SR in the Δ(1–209) mutation rescues this defect by unleashing the LH, suggesting that the SR region is not essential for vRNP formation. The SR region not only contributes to RNA binding, but also has been shown to cooperatively enhance the RNA binding affinity of the NTD and CTD (27,49–51). We therefore speculate that by increasing the RNA binding activity of the NTD, the SR may improve the NTD’s ability to act as a regulatory switch and release the LH.

Our evidence suggests that phosphorylation of the SR region inhibits vRNP formation not simply by weakening RNA binding to the SR region, but also by promoting an inhibitory interaction with the RNA-binding site at the CTD. Our results are consistent with previous studies showing that phosphorylated N protein cannot form compact vRNPs for packaging, instead maintaining RNA in an elongated, uncompacted state (25,26). Phosphorylation inhibits the assembly of VLPs but also appears to be important for efficient viral genome replication, revealing a trade-off between the opposing effects of phosphorylation (61). The Δ(1–209) mutation has been implicated as an evolutionary strategy to resolve the adaptive conflict between replication and assembly. By removing the SR region and its associated phosphorylation sites, Δ(1–209) may provide a fitness benefit by complementing highly phosphorylated full-length N protein with kinase-independent assembly by truncated N (61).

The processes of N protein-mediated RNA packaging and transcription represent important targets for viral inhibition. The development of strategies to reduce viral infectivity requires a molecular understanding of the viral replication cycle and the specific architecture and regulation of viral packaging. The LH has been shown to interact with Nsp3, a transmembrane protein localized to double-membrane vesicles at the RTC (17,74–76). Therefore, in addition to driving vRNP formation by promoting N self-association, the LH may also mediate N protein’s interaction with the RTC. Given the high conservation of the disordered central linker region in betacoronaviruses, inhibition of the LH represents a promising target for novel coronavirus antiviral therapies.

### Experimental Procedures

#### N protein expression and purification

Full-length and mutant N proteins were prepared as described previously (25,26). The ‘wild-type’ N protein used in these studies is from the 2019 WH-Human 1 strain. Briefly, codon-optimized synthetic DNAs encoding N protein were ordered as gBlocks from Integrated DNA Technologies (IDT) and inserted into pET28a expression vectors by Gibson assembly. A 6xHis-SUMO tag was added to the N terminus of all N protein constructs. N protein mutants were constructed by site-directed mutagenesis and Gibson assembly. N protein vectors were transformed into *E. coli* BL21 Star (Thermo #C601003) for expression and grown in TB+Kanamycin to OD_600_ = 0.6 at 37°C. Protein expression was induced with 0.4 mM IPTG at 16°C overnight. Cells were harvested, washed with PBS, snap frozen in LN_2_, and stored at −80°C until lysis. N protein was purified under denaturing conditions to remove nucleic acids. Thawed cells were resuspended in buffer A (50 mM HEPES pH 7.5, 500 mM NaCl, 10% glycerol, and 6 M urea) and lysed by sonication. Lysed cells were centrifuged at 30,000 rpm for 30 min to remove cell debris, and clarified lysate was bound to Ni-NTA agarose beads (QIAGEN #30230) for 45 min at 4°C. Ni-NTA beads were washed three times with ten bed volumes of buffer A and eluted with buffer B (50 mM HEPES pH 7.5, 500 mM NaCl, 10% glycerol, 250 mM imidazole, and 6 M urea). The eluate was concentrated in centrifugal concentrators (Millipore Sigma #UFC803024), transferred to dialysis tubing (Spectrum Labs #132676), and renatured by overnight dialysis in buffer C (50 mM HEPES pH 7.5, 500 mM NaCl, 10% glycerol). Recombinant Ulp1 catalytic domain (expressed and purified separately from *E. coli*) was added to renatured proteins to cleave the 6xHis-SUMO tag. Cleaved protein was injected onto a Superdex 200 10/300 size-exclusion column equilibrated in buffer C. Peak fractions were pooled, concentrated, frozen in LN_2_, and stored at −80°C.

#### RNA preparation

The template for *in vitro* transcription of 5’-600 RNA was ordered as a gBlock from IDT and inserted by Gibson assembly into a pUC18 vector with a 5’ T7 promoter sequence. An insert containing the 5’-600 DNA plus the 5’ T7 sequence was excised by EcoR1 digestion and purified by size-exclusion chromatography on a Sephacryl 1000 column equilibrated in TE buffer (10 mM Tris pH 8, 1 mM EDTA). Peak fractions of the purified DNA insert were pooled and stored at −20°C. The SL8 template was generated by PCR of synthetic DNA (IDT) to contain a 5’ T7 transcription start site. PCR-amplified DNA was purified and concentrated by spin column (Zymo Research #D4004) before being used to generate RNA.

All RNA synthesis was performed using the HiScribe T7 High Yield RNA synthesis kit (NEB E2040S) according to the manufacturer’s protocol. Following incubation at 37°C for 3 h, *in vitro* synthesized RNA was purified and concentrated by spin column (Zymo Research #R1018). To promote formation of proper RNA secondary structure, all purified RNAs were heat denatured at 95°C for 2 min in a preheated metal heat block and then removed from heat and allowed to cool slowly to room temperature over the course of ∼1 h. RNA concentration (A260) was quantified by nanodrop.

#### Preparation of ribonucleoprotein complexes

The day before each experiment, N protein was dialyzed into reaction buffer (25 mM HEPES pH 7.5, 70 mM KCl) overnight. The day of analysis, RNA was heat-denatured and cooled slowly to allow for proper secondary structure formation. To assemble vRNP complexes, 256 ng/μl RNA was mixed with 15 μM N protein in a total volume of 10 μl and incubated for 10 min at 25°C. RNA concentration was 256 ng/μl, regardless of RNA length, to ensure that all mixtures contained the same nucleotide concentration. Samples containing SL8 stem-loop RNA were crosslinked by addition of 0.1% glutaraldehyde for 10 min at 25°C and then quenched with 100 mM Tris pH 7.5. Samples containing 5’-600 RNA were not crosslinked. After assembly, vRNP complexes were analyzed by gel electrophoresis.

#### RNA gel electrophoresis

After assembly, the 10 μl vRNP mixtures were diluted 1:10 in dilution buffer (25 mM HEPES pH 7.5, 70 mM KCl, and 10% glycerol). Diluted vRNP mixture (2 μl) was loaded onto a 5% polyacrylamide native TBE gel (Bio-Rad) and run at 125 V for 80 min at 4°C. Another aliquot of diluted sample (1 μl) was denatured by addition of 4 M urea and Proteinase K (40 U/ml; New England Biolabs #P8107S), incubated for 5 min at 65°C, loaded onto a 6% polyacrylamide TBE-Urea Gel (Thermo Fisher), and run at 160 V for 50 min at room temperature. Gels were stained with SYBR Gold (Invitrogen) and imaged on a Typhoon FLA9500 Multimode imager set to detect Cy3.

#### Glycerol gradient centrifugation

Glycerol gradients were assembled as previously described (69), with slight modifications. Briefly, 10 to 40% glycerol gradients (dialysis buffer containing 10% or 40% glycerol) were poured and mixed with the Gradient Master (BioComp). Fresh 0.1% glutaraldehyde was added to the 40% glycerol buffer prior to gradient assembly. vRNP samples (140 μl of 15 μM N with 256 ng/μl RNA) were gently added on top of the assembled 5 ml gradients, and samples were centrifuged in a prechilled Ti55 rotor at 35,000 rpm for 17 h. Gradient fractions were collected by puncturing the bottom of the tube with a butterfly needle and collecting two drops per well. For analysis by negative stain electron microscopy and mass photometry, indicated fractions were combined, and buffer exchanged using centrifugal concentrators (Millipore Sigma #UFC510024). Concentrated samples were then re-diluted 1:10 with dialysis buffer (0% glycerol) and re-concentrated. Samples were diluted and re-concentrated three times.

#### Mass photometry

Mass photometry experiments were performed using a OneMP instrument (Refeyn). A silicone gasket well sheet (Grace Bio-Labs) was placed on top of a microscope coverslip and positioned on the microscope stage. Reaction buffer (10 μl) (25 mM HEPES pH 7.5, 70 mM KCl) was first loaded into the well to focus the objective, after which 1 μl of vRNP complex sample was added to the reaction buffer, mixed, and measured immediately. Crosslinked samples containing SL8 RNA were diluted 1:10 before a second 1:10 dilution directly on the coverslip.

The mass photometer was calibrated with NativeMark Unstained Protein Standard (Thermo #LC0725). Mass photometry data were acquired with AcquireMP and analyzed with DiscoverMP software (Refeyn). Mass photometry data are shown as histograms of individual mass measurements. Peaks were fitted with Gaussian curves to determine the average molecular mass of the selected distributions. Each condition was independently measured at least twice.

#### Negative stain electron microscopy

2.5 μl of vRNP samples were applied to a glow-discharged Cu grid covered with continuous carbon film and stained with 0.75% (w/v) uranyl formate. A Tecnai T12 microscope (ThermoFisher FEI Company), operated at 120 kV, was employed to analyze these negatively stained grids. Micrographs were recorded at a nominal magnification of 52,000× using a Gatan Rio 16 camera, corresponding to a pixel size of 1.34 Å on the specimen. All images were analyzed using ImageJ. Micrographs were labeled with a corresponding scale bar of 20 nm, and particles were manually extracted individually using cropped boxes for illustration. 2D averaging of particles was not performed due to their heterogeneity.

#### Fluorescence Polarization

Fluorescence was measured on a K2 Multifrequency Fluorometer. Fluorescent N_10_ RNA (a 10-nt degenerate sequence, tagged with a 3’-FAM fluorescent dye) was ordered from IDT. Undialyzed N protein constructs were diluted 1:10 in reaction buffer (25 mM HEPES pH 7.5, 70 mM KCl) and concentrated using centrifugal concentrators (Millipore Sigma #UFC510024). 10 nM fluorescent RNA was mixed with 10 μM N protein in 90 μl reaction buffer, added to a 3×3 mm quartz cuvette (Hellma #1052518540), and incubated for 5 min. Protein concentration was gradually reduced by serial dilutions with a solution of reaction buffer containing 10 nM N_10_ RNA to keep fluorescent RNA concentration constant. The volume in the cuvette was maintained throughout all titration experiments by adding 90 μl dilution buffer to the sample cuvette, mixing by pipetting, and removing an equal volume of solution. RNA was excited with polarized light at 488 nm and emission was recorded at 520 nm. Data from three independent N protein titrations were fit to a one-site binding curve using GraphPad Prism to determine K_D_.

N protein was phosphorylated *in vitro* for fluorescence polarization analysis. Protein kinases were purchased from Promega (SRPK1: #VA7558, GSK-3β: #V1991, CK1ε: V4160). 15 μM Δ(1–175) N protein and 80 nM SRPK, GSK3, and CK1 were mixed in 100 μl kinase reaction buffer (25 mM HEPES pH 7.5, 35 mM KCl, 10 mM MgCl_2_, 1 mM DTT, 0.5 mM ATP). The reaction was incubated for 30 min at 30°C and quenched with 5 mM EDTA. A separate control reaction lacking ATP was prepared in the same way. Both reactions were diluted 1:10 in reaction buffer (25 mM HEPES pH 7.5, 70 mM KCl) and concentrated using centrifugal concentrators (Millipore Sigma #UFC510024). 10 nM fluorescent RNA was mixed with 10 μM N protein in 90 μl reaction buffer, added to a 3×3 mm quartz cuvette (Hellma #1052518540), and incubated for 5 min. Protein concentration was gradually reduced by serial dilutions, while keeping RNA concentration and cuvette volume constant. RNA was excited with polarized light at 488 nm and emission was recorded at 520 nm. Data from two independent N protein titrations were fit to a one-site binding curve using GraphPad Prism to determine K_D_.

#### Virus-like particle (VLP) Assay

VLPs were produced and analyzed as previously described (61,63). 293T cells were plated at 50,000 cells/well in 150 μl of DMEM containing 10% FBS and 1% penicillin/streptomycin in 96 well plates and incubated for 24 h. 1.5 μg of plasmids encoding N protein (0.33), Firefly luciferase with packaging signal T20 (0.5), Spike (0.0033), and M-IRES-E (0.165) at indicated mass ratios were diluted in 75 μl room temperature Opti-MEM. 4.5 μl of 1 mg/ml polyethylenimine (pH 7.0) was diluted in 75 μl Opti-MEM and used to resuspend plasmid dilutions. The diluted plasmid transfection mix was incubated at room temperature for 20 min. For transfection into 96 well plates, 18.75 μl of resulting transfection mix was added to each well and mixed by pipetting up and down carefully. For GSK3 inhibition, 17 μl of 12.5 μM CHIR98014 in media was added to wells immediately after adding the transfection mix. Control wells were treated with 17 μl of additional media. 2 days after transfection, 65 μl of supernatant was filtered using a 96-well filter plate (Pall 0.45 μm AcroPrep™ Advance Plate, Short Tip filter plates, polyethersulfone) by centrifuging the filter plate at 100xg for 2 min. 50 μl of 1×10^6^/ml 293T-ACE2/TMPRSS2 cells was added to the VLP-containing filtrate and incubated for 24 h. The supernatant was removed and 20 μl of Promega passive lysis buffer (1X) was added to the cells. The cells were then incubated on a shaker for 20 minutes. Using a Tecan Spark, 30 μl of luciferase assay buffer was dispensed into each well and mixed for 30 s by shaking. Luminescence was measured using the auto attenuation function and 1000 ms integration time.

## Data availability

All data are included in the article, or available from the corresponding author D.O.M.

## Supporting information

This article contains supporting information.

## Conflict of interest

The authors declare that they have no conflicts of interest with the contents of this article.

## Acknowledgments

We thank Henry Ng and Emmy Delaney for discussions, technical assistance, and comments on the manuscript, and Hayden Saunders and Arthur Charles-Orszag for technical assistance.

## Funding and additional information

This work was supported by grants (to D.O.M.) from the National Institute of General Medical Sciences (R35-GM118053) and the UCSF Program for Breakthrough Biomedical Research, which is funded in part by the Sandler Foundation. Funding was also provided by the National Institutes of Allergy and Infectious Diseases (R21-AI159666 to J.A.D.), the Howard Hughes Medical Institute (Y.C. and J.A.D.) and the J. David Gladstone Institutes (J.A.D.).

## Supporting Information

**Figure S1.**
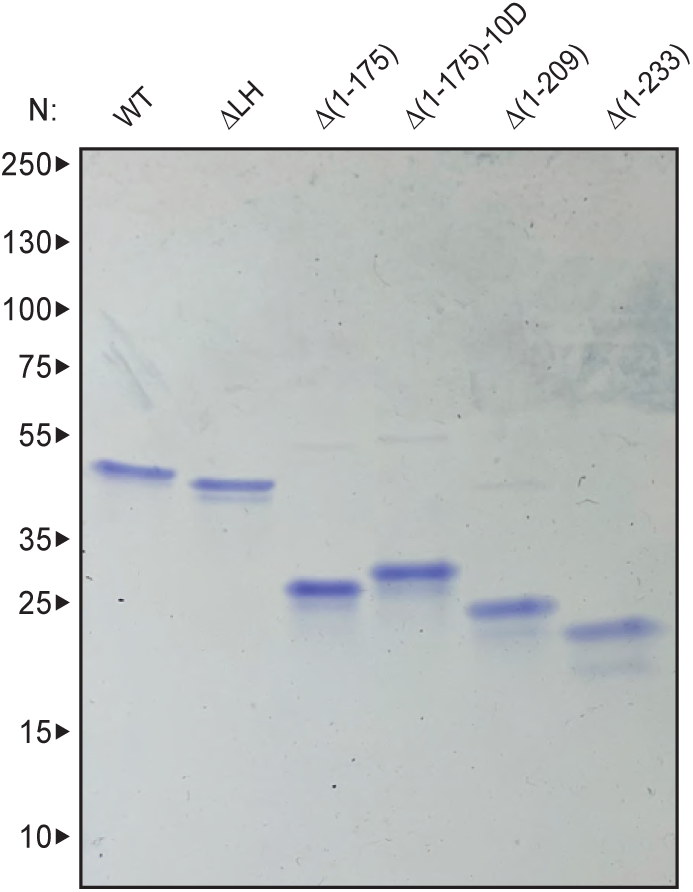
Analysis of N protein deletion mutants. SDS-PAGE of N protein constructs used in this study, stained with Coomassie Blue. Molecular weight markers shown on the left (kDa).

**Figure S2.**
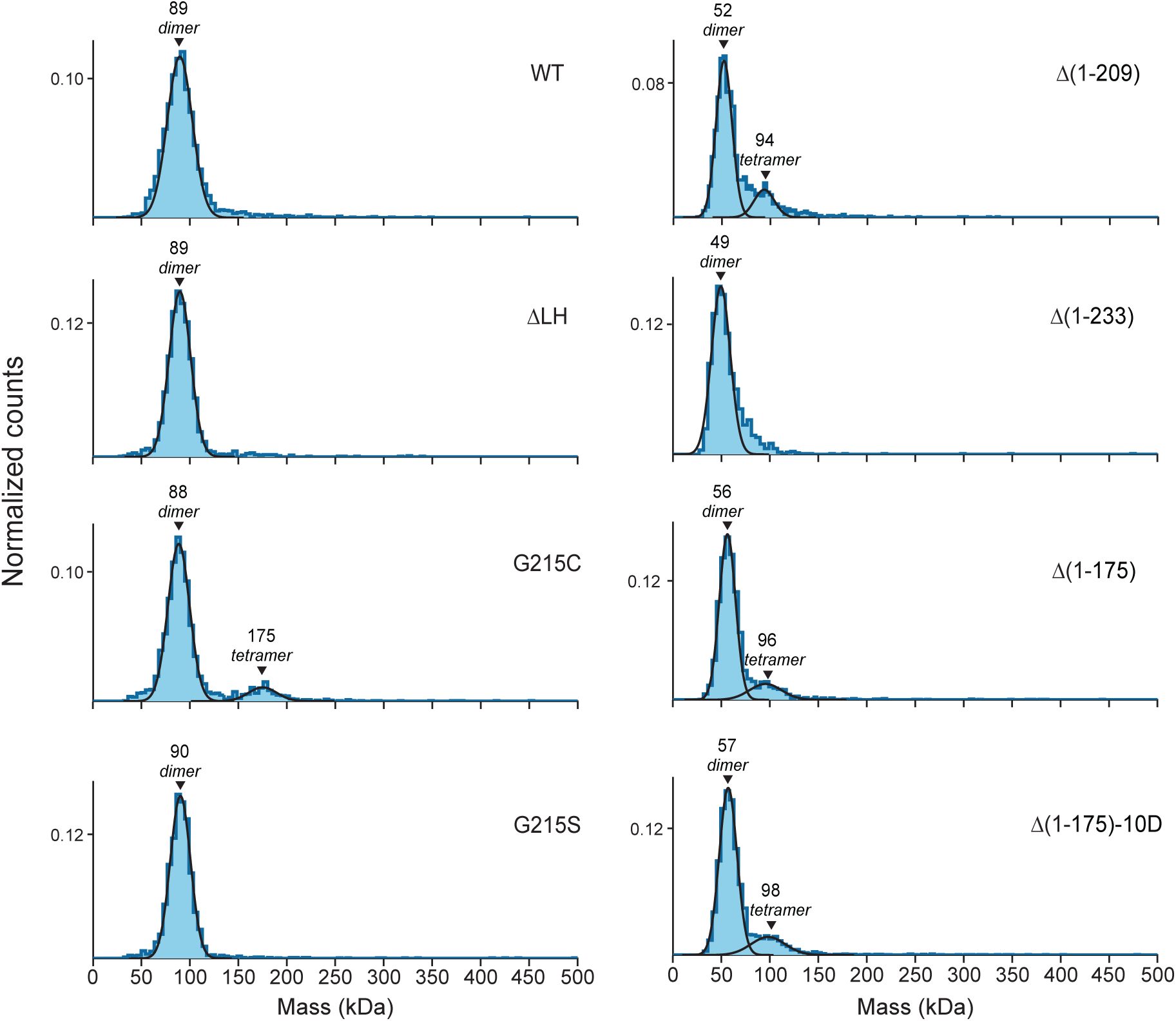
Oligomerization of uncrosslinked N protein mutants. Mass photometry analysis of indicated N protein mutants in the absence of crosslinker, analyzed in parallel to the crosslinked preparations shown in Fig. 2. Data were fit to Gaussian distributions, with mean molecular mass and corresponding oligomeric state indicated above each peak. Representative of two independent experiments (Table S1). LH, leucine helix; N, nucleocapsid; WT, wild type.

**Table S1.**
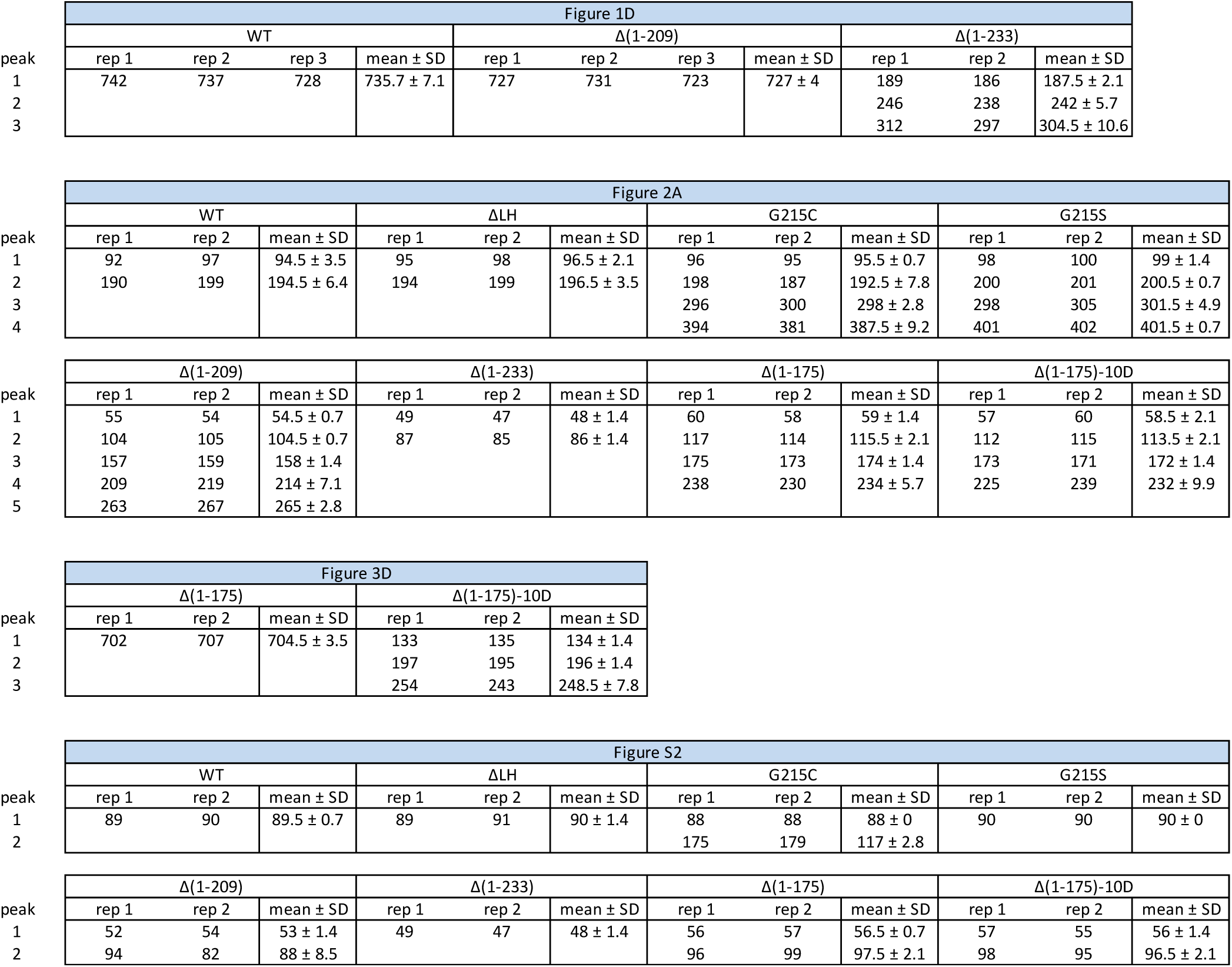
Summary of mass photometry results (kDa).

## Notes

### Competing Interest Statement

The authors have declared no competing interest.

